# p16-3MR: a novel model to study cellular senescence in cigarette smoke-induced lung injuries during aging

**DOI:** 10.1101/2021.02.23.432412

**Authors:** Gagandeep Kaur, Isaac K. Sundar, Irfan Rahman

## Abstract

Cellular senescence and lung aging are associated with the pathogenesis of Chronic Obstructive Pulmonary Disease (COPD). COPD progresses with aging, and chronic smoking is the key susceptibility factor in lung pathological changes concurrent with biological aging. However, these processes involving cigarette smoke (CS)-mediated lung cellular senescence are difficult to distinguish. One of the impediments to study cellular senescence in relation to age-related lung pathologies is the lack of a suitable *in vivo* model. In view of this, we provide evidence that supports the suitability of p16-3MR mice to study cellular senescence in CS-mediated and age-related lung pathologies. p16-3MR mice has a trimodal reporter fused to the promoter of p16^INK4a^ gene that enables detection, isolation and selective elimination of senescent cells, thus making it a suitable model to study cellular senescence. To determine its suitability in CS-mediated lung pathologies, we exposed young (12-14 months) and old (17-20 months) p16-3MR mice to 30-day CS exposure and studied the expression of senescent genes (p16, p21 and p53) and SASP-associated markers (MMP9, MMP12, PAI-1, and FN-1) in air- and CS-exposed mouse lungs. Our results showed that this model could detect cellular senescence using luminescence and isolate cells undergoing senescence with the help of tissue fluorescence in CS-challenged young and old mice. Our results from the expression of senescence markers and SASP-associated genes in CS-challenged young and old p16-3MR mice were comparable with increased lung cellular senescence and SASP in COPD. We further showed age-dependent alteration in the (i) tissue luminescence and fluorescence, (ii) mRNA and protein expressions of senescent markers and SASP genes, and (iii) SA-β-gal activity in CS-challenged young and old p16-3MR mice as compared to their air controls. Overall, we showed that p16-3MR is a competent model to study cellular senescence in age-related pathologies and could help understand the pathobiology of cellular senescence in lung conditions like COPD and fibrosis.

## INTRODUCTION

Cigarette smoke (CS) can cause DNA damage and cellular senescence, leading to premature lung aging ^1–4^. Reports prove a critical role of cigarette smoke-induced cellular senescence in the development of Chronic Obstructive Pulmonary disease (COPD) or emphysema ^3,5–7^. Markers of senescence, including p16 and p21, are shown to be upregulated in both the airway epithelium and endothelium of lung specimens from patients with COPD, thus proving that cellular senescence has an important role in the pathophysiology of COPD ^3,6,8^. Cellular senescence refers to the state of irreversible cell cycle arrest in somatic cells in response to intrinsic stressors (DNA damage) or extrinsic stressors (oxidative stress) ^9^. Unlike cells that have undergone apoptosis, senesced cells remain metabolically active and continue to affect their surrounding cells after having undergone specific phenotypic changes themselves. One of these phenotypic changes is an increased production of extracellular vesicles and an increased secretion of inflammatory cytokines and interleukins. Changes in the secretory profile of a senescent cell reshapes not only its microenvironment, but also that of its surrounding cells, and is termed as senescence-associated secretory phenotype (SASP) ^9,10^. SASP has been shown to be linked to chronic inflammation, which is a ubiquitous component of aging tissues and most age-related diseases including COPD and Idiopathic pulmonary fibrosis (IPF) ^10–12^. Mounting evidence proves that the elimination of these senescent cells can prevent the development and/or exacerbation of certain age-related pathologies ^13–17^.

The major impediment in studying the role of senescent cells in age-related pathologies is the lack of a suitable reporter model. To overcome this limitation, Demaria et al. created a novel mouse model p16-3MR which can (a) detect senescent cells in living animals, (b) enable the isolation of senesced cells from mouse tissues, and (c) eliminate senescent cells upon treatment with a drug otherwise ineffective in wild-type mice. p16-3MR mice carries a transgene consisting of a trimodal reporter under the regulation of a p16 promoter ^18^. p16 is a tumour suppressor gene involved in regulating cellular senescence, specifically through its induction of G1 cell cycle arrest by inhibiting cyclin-dependent kinases ^3,19^. Theoretically, p16-3MR is a useful model for studying cellular senescence in various age-related pathologies ^15–17^; however, to our knowledge, this model's efficacy in studying lung pathologies induced by cigarette smoke exposure has not been tested.

Considering this, we studied the suitability of using p16-3MR as a model to study cellular senescence in cigarette smoke-induced pulmonary pathologies. Previous reports by our group have shown that chronic exposure to cigarette smoke induced the upregulation of cellular senescence as well as the expression of p16 in C57BL/6J mice, independent of age ^20^. In this study, we determined whether p16-3MR reporter mice could be used to study the role of lung cellular senescence in the pathophysiology of COPD. We analysed the changes in the expression of SASP markers following 30-day CS exposure in both young and old p16-3MR mice. Using In Vivo Imaging System (IVIS) and fluorescence microscopy of lung tissues we show that p16-3MR mice can be successfully used to visualize and isolate senescent cells from the lungs of CS-exposed mice.

## RESULTS

### Sub-chronic CS exposure augments luminescence indicative of p16 expression in the lungs of p16-3MR reporter mice

p16-3MR mice are unique model designed to study cellular senescence by the fusion of senescence-sensitive promoter of p16^INK4a^ and its adjacent gene p19^Arf^ with a trimodal reporter constituting of functional domains for LUC (Renilla Luciferase), mRFP (monomeric red fluorescent protein) and HSV-TK (truncated herpes simplex virus 1 thymidine kinase). We tested the suitability of this model to study CS-induced cellular senescence by exposing young and old p16-3MR mice to sub-chronic CS and harvested the lung tissues. The 3MR-expressing cells were first detected using luminescence in the lung tissues from air- and CS-exposed young and old mice. Our results demonstrated a significant increase in the lung tissue luminescence following sub-chronic CS exposure (30 days) in young p16-3MR mice as compared to their older counterparts. However, the augmentation in cellular senescence, as indicated by the increase in tissue luminescence, was not significant in old p16-3MR mice following 30-day CS-challenge (**Figure 1**).

**Figure 1:**
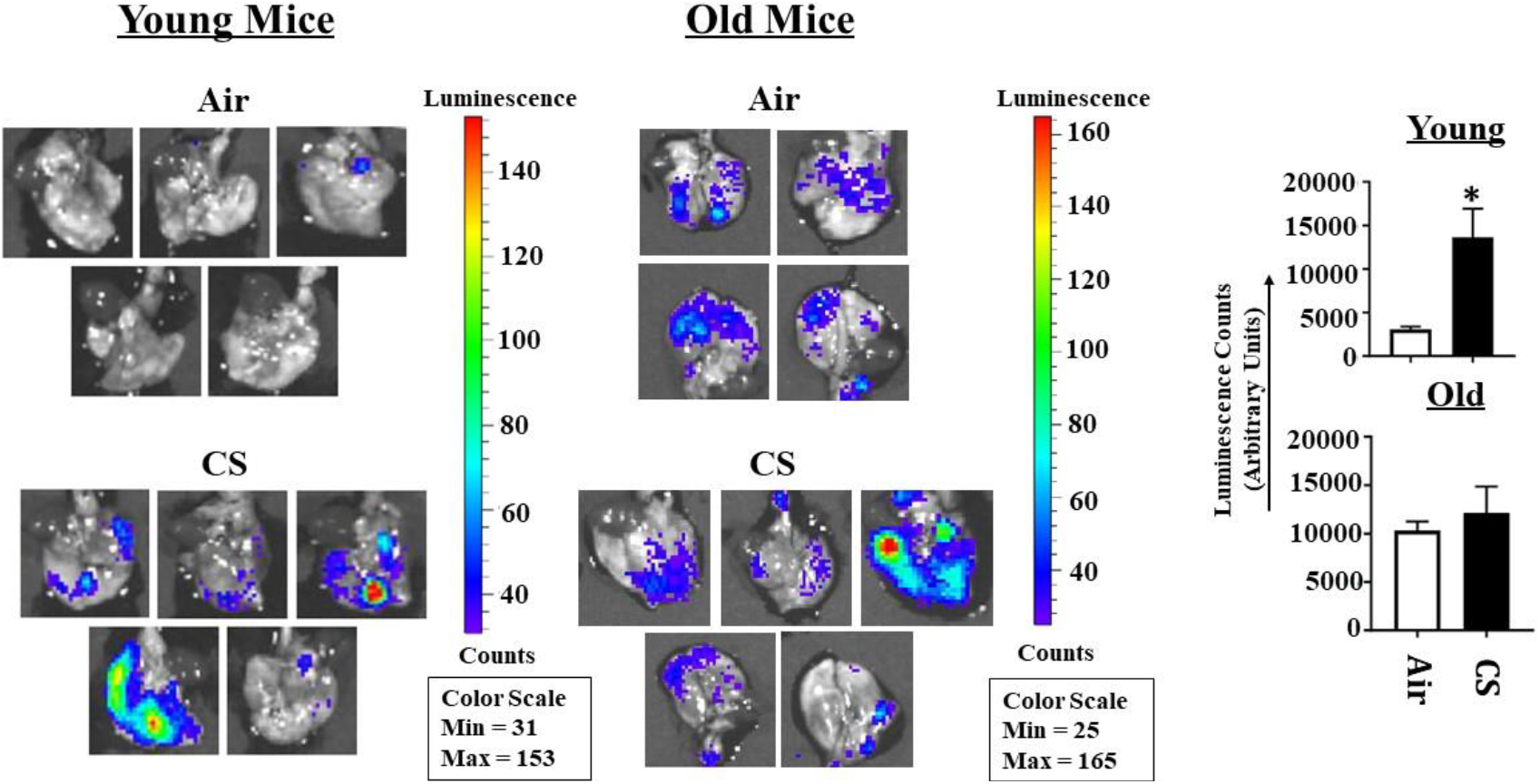
Detection of cellular senescence using tissue luminescence in the lungs of CS-exposed p16-3MR mice. Young and old p16-3MR mice were subjected to sub-chronic (30 days) CS exposure. The lung tissues from air- and CS-exposed mice were harvested and the lung tissue luminescence was measured using IVIS® Spectrum multispectral imaging instrument. Data was normalized using the tissue luminescence from the lung tissues of wild-type (C57BL/6J) mice. Representative images of tissue luminescence from each sample were provided and the quantified luminescence counts from each sample plotted alongside. Data are shown as mean ± SEM (n = 4–5/group). * p < 0.05; vs Air as per Student’s t-test for pairwise comparisons.

### Sub-chronic exposure to CS results in a significant increase in the mRFP expression in the lungs of young p16-3MR mice

To sort the tissues/cells undergoing cellular senescence, we tested the mRFP fluorescence and expression in the lung tissues from both CS-challenged young and old p16-3MR mice. We first employed IVIS imaging to measure the mRFP fluorescence in air- and CS-exposed lungs from younger and older groups of p16-3MR mice. Our results showed a significant increase in the lung tissue fluorescence (mRFP) on CS-challenge in both age groups, but we observed a lot of background signal in our samples (**Figure 2**). We suspect that this auto-fluorescence was due to the possibility of CS-induced tar deposition in the lung during sub-chronic CS exposure. Thus, we measured the tissue fluorescence in the lung tissue sections from air/CS-exposed young and old p16-3MR mice. Our investigations showed a pronounced upregulation in mRFP expression of CS-challenged young p16-3MR mice relative to their respective age-related air controls. Contrarily, the increase in the mRFP expression in the lungs of CS-challenged old p16-3MR was not significant (**Figure 3a**). Furthermore, mRNA expression analyses for mRFP expression correlated with our previous findings, further substantiating our results (**Figure 3b**).

**Figure 2:**
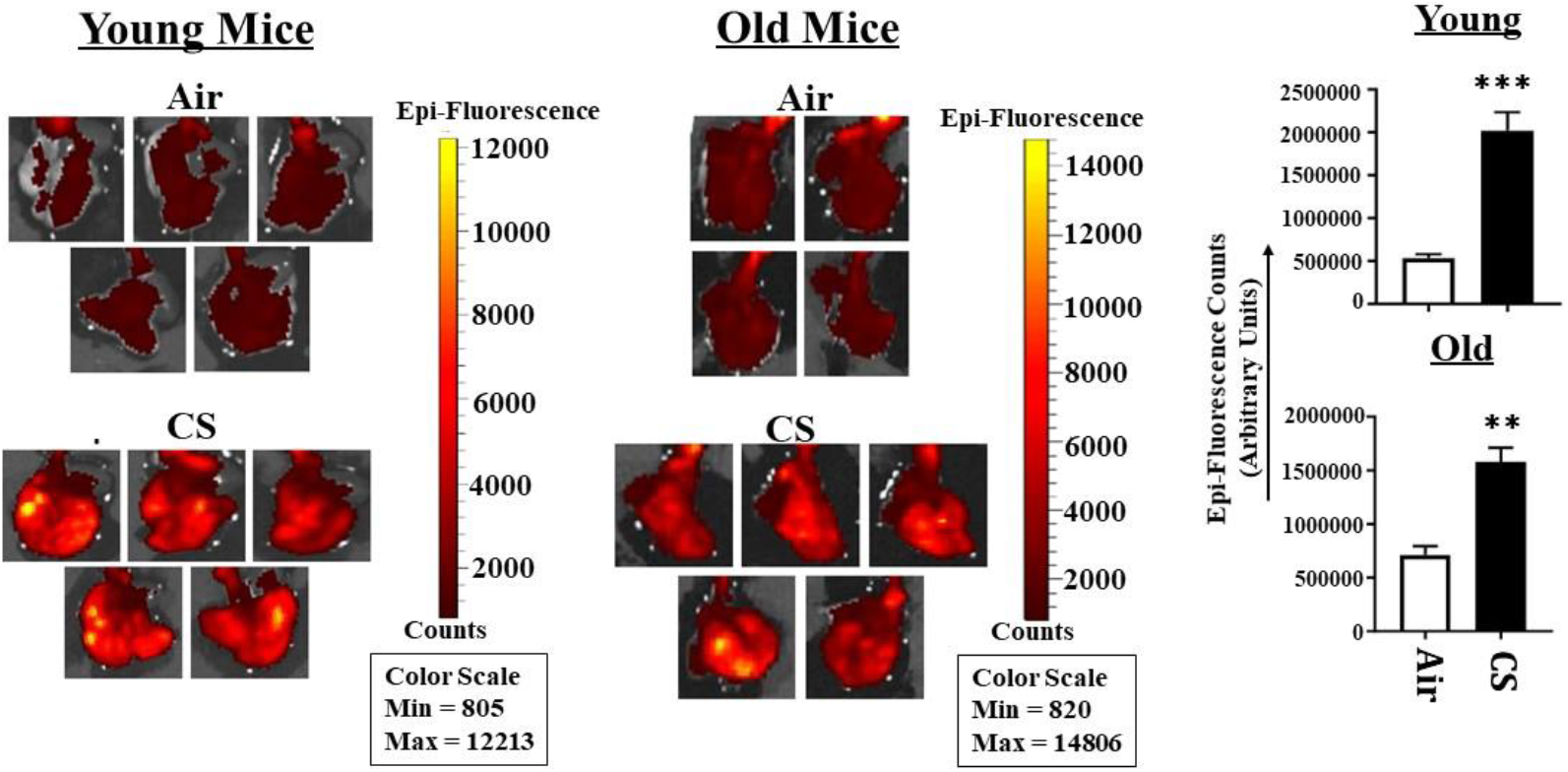
Tissue fluorescence in the lungs of CS-exposed young and old p16-3MR mice using IVIS imaging. Young and old p16-3MR mice were subjected to sub-chronic (30 days) CS exposure. The lung tissues from air- and CS-exposed mice were harvested and the lung tissue fluorescence was measured using IVIS® Spectrum multispectral imaging instrument at excitation and emission maximum of 535 and 580 nm respectively. Data was normalized using the tissue luminescence from the lung tissues of wild-type (C57BL/6J) mice. Representative images of n = 4–5/group were provided and the quantified fluorescence counts plotted as mean ± SEM. \*\**p* < 0.01, \*\*\**p* < 0.001; vs Air as per Student’s t-test for pairwise comparisons.

**Figure 3:**
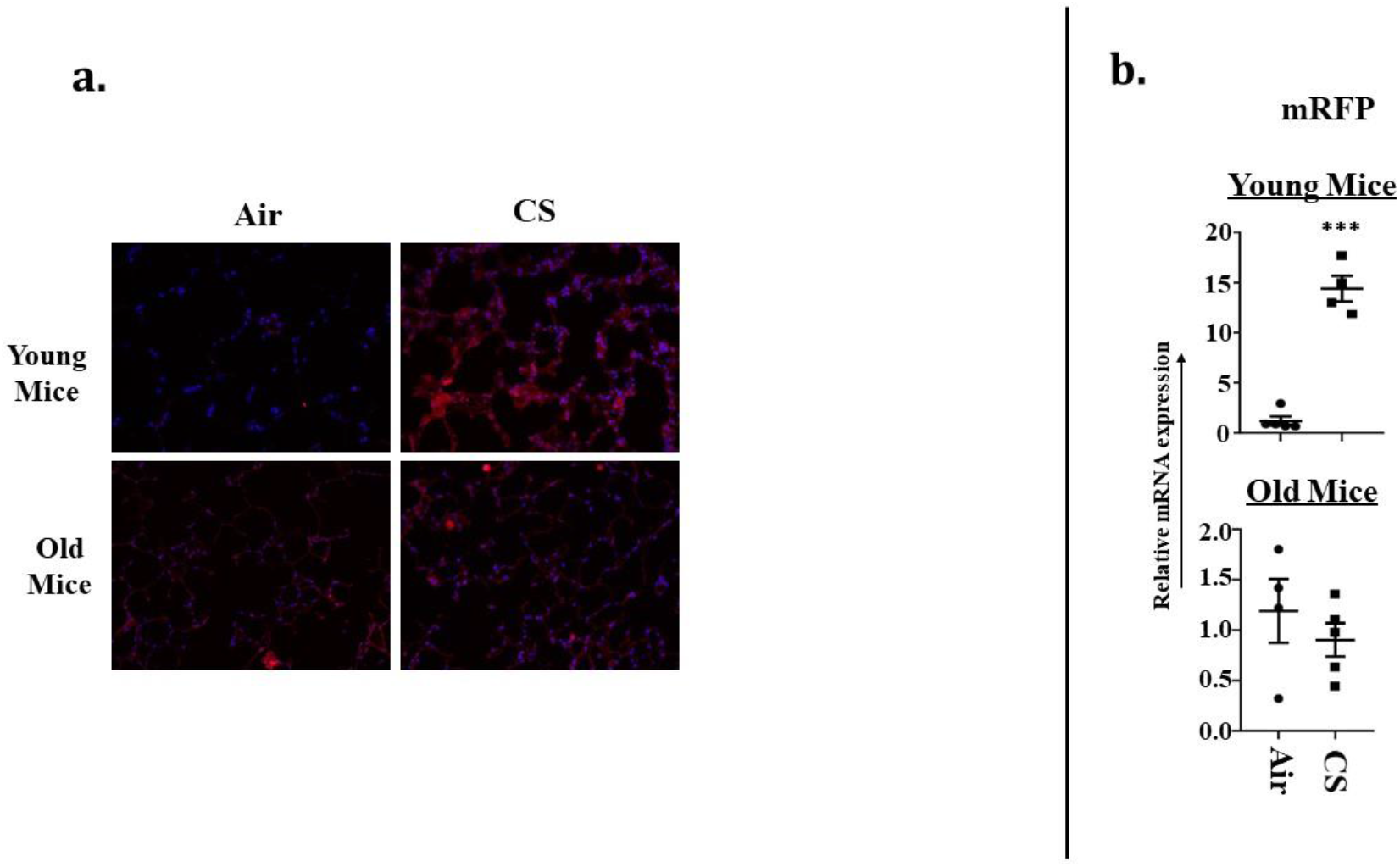
Senescent cells observed in lung tissues from CS-exposed p16-3MR mice by fluorescence microscopy. (a) Lung tissue sections from air and CS-exposed younger and older p16-3MR mice were stained with DAPI and tissue fluorescence (mRFP) was observed using Nikon Eclipse Ni-U microscope. Representative images of n=4–5/group were provided. Original magnification, x200. (b) mRNA expression of mRFP in the lung tissue was examined by qPCR. Bar graphs represent the mean normalized fold change of respective air- vs. CS-exposed mouse lungs using ∆∆Ct method. Data are shown as mean + SEM (n = 4–5/group). ***p < 0.001, vs Air as per Student’s t-test for pairwise comparisons.

### Increased expression of cellular-senescence markers (p16 and p21) following sub-chronic CS exposure in young p16-3MR mice

To further confirm our findings, we next determined the expression of the markers of cellular senescence p16 and p21 in our study model at both transcriptional and translational levels. The mRNA levels of senescence markers p16^INK4a^ and p21 were significantly upregulated in young CS-exposed p16-3MR mice, confirming the presence of senescent cells following sub-chronic CS exposure (**Figure 4a**). Immunoblotting results supported the findings from the aforementioned mRNA expression, further substantiating our claims (**Figure 4b**). Contrarily, as expected, we did not find a significant increase in mRNA or protein expression of p16^INK4a^ in older CS-challenged mice as compared to their age-related air controls (**Figure 4a&b**). In fact, the protein expression of p16 was reduced on CS-exposure in older p16-3MR mice **(Figure 3b).** This finding was not surprising due to the baseline expression of p16^INK4a^ gene for the air-controls being elevated in the older mice, thus conferring with the normal age-related process. Interestingly, despite observing a significant increase in the mRNA expression of the p21 gene in both young and old CS-exposed mice, we demonstrated a marked decrease in its protein expression in older CS-exposed mice relative to their respective air control (**Figure 4a&b**).

**Figure 4:**
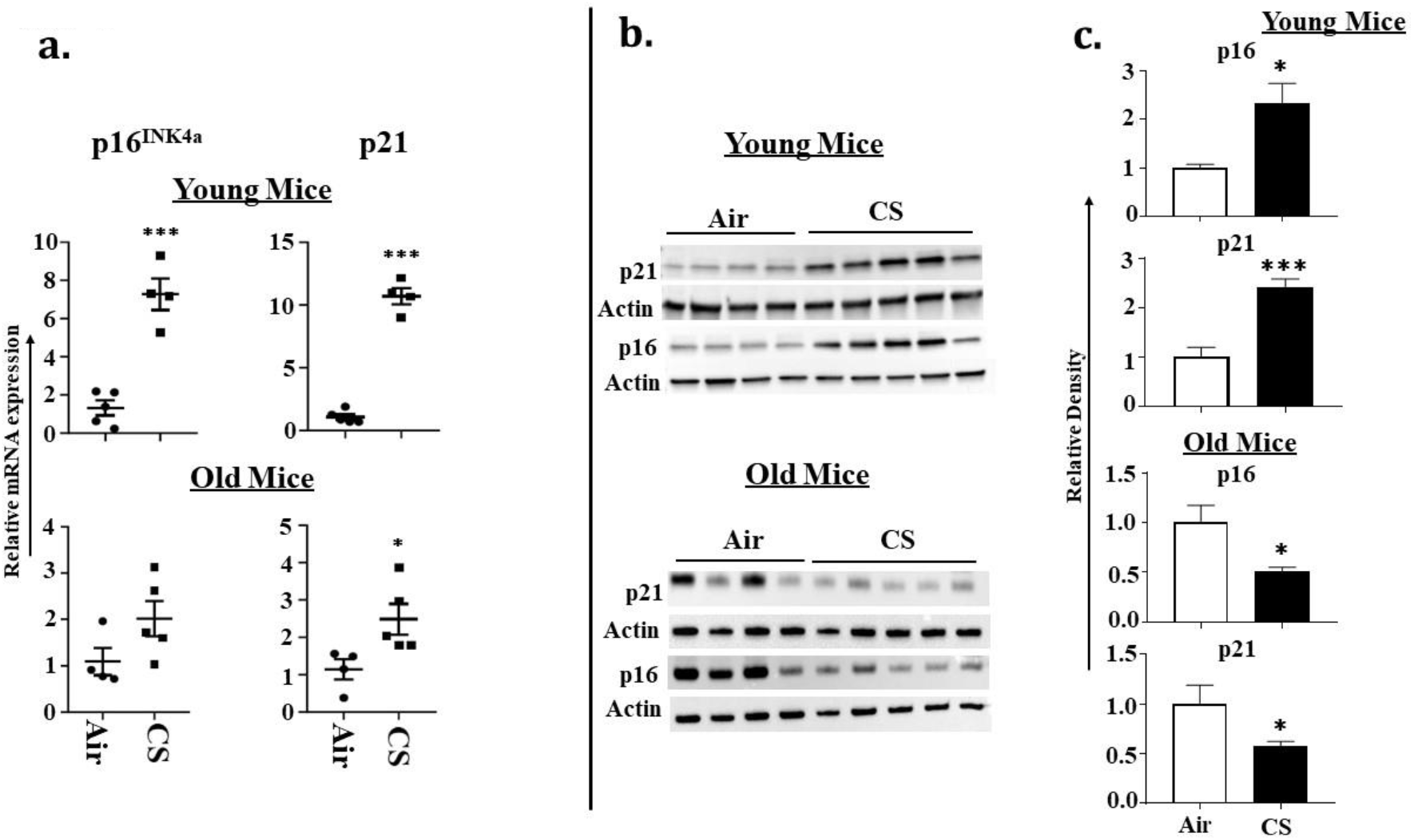
Increased transcriptional and translational expression of senescence markers in CS-exposed p16-3MR mice. Younger and older mice were exposed to sub-chronic CS for 30 days. (a) mRNA and (b) protein expressions of early- (p21) and late-stage (p16) senescence markers were measured in the lungs homogenates using qPCR and immunoblotting analyses respectively. β-actin/Tubulin was used as loading controls. (c)The band intensity was measured by densitometry and data were shown as fold change relative to respective control group. Full Gels/blots with bands (unedited/uncropped electrophoretic gels/blots) obtained from air- and CS-exposed younger and older mouse samples from the same experiments were processed in parallel are shown (see Supplementary information as Suppl Figs. 3-4). Data are shown as mean ± SEM (n = 4–5/group). \**p* < 0.05, \*\*\**p* < 0.001 vs Air as per Student’s t-test for pairwise comparisons.

### Age-dependent changes in the mRNA and protein expression of SASP-associated genes in CS exposed p16-3MR mice

To determine the age-related changes in the protein expression of various SASP-associated proteins, we studied the expression of MMP12, MMP9, FN-1, PAI-1 and p53 in air- and CS-exposed younger and older p16-3MR mice. Consistent with the gene expression studies performed by our groups, we found a significant age-independent increase in the mRNA expression of MMP12 (**Figure 5**). However, the protein expression data did not show any notable variation in the expression of MMP12 in the lung of air- or CS-exposed young and old p16-3MR groups. In addition, no significant change in the expression of MMP9 was observed in response to sub-chronic CS exposure in young and old groups of p16-3MR mice, thus suggesting no changes in the extracellular matrix (ECM) composition on CS (30 day) exposure (**Figure 5**).

**Figure 5:**
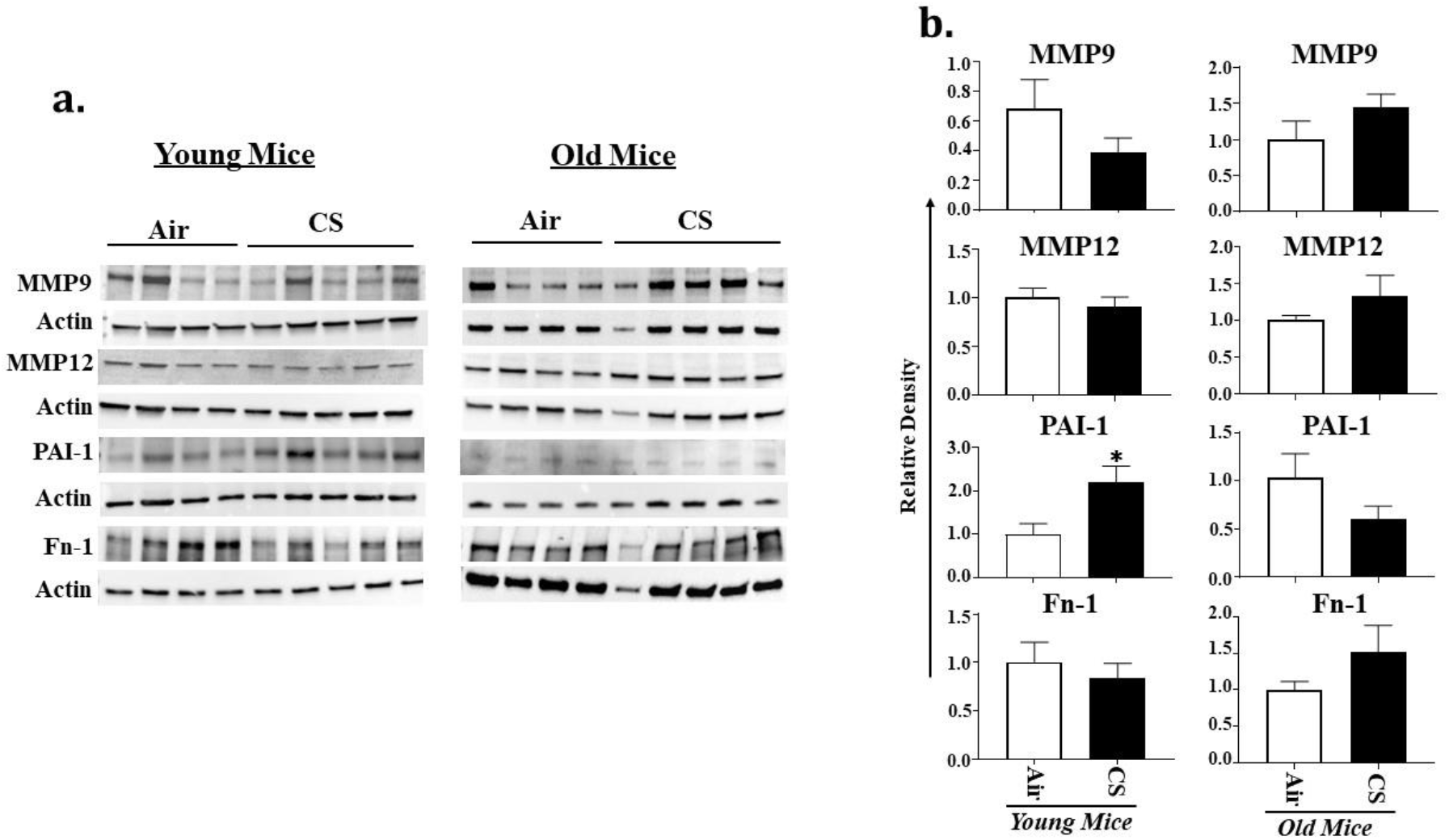
Altered protein abundance of SASP-associated markers in CS-exposed p16-3MR mice. (a) Younger and older p16-3MR mice were exposed to sub-chronic CS for 30 days and the expression of SASP-associated markers MMP9, MMP12, PAI-1 and FN-1-were determined using immunoblotting analyses. β-actin was used as loading controls. (b) The band intensity was measured by densitometry and data were shown as fold change relative to respective control group. Full Gels/blots with bands (unedited/uncropped electrophoretic gels/blots) obtained from air- and CS-exposed younger and older mouse samples from the same experiments were processed in parallel are shown (see Supplementary information as Suppl Figs. 5-8). Data are shown as mean ± SEM (n = 4–5/group). \**p* < 0.05 vs Air as per Student’s t-test for pairwise comparisons.

We further demonstrated a pronounced upregulation in the expression of PAI-1, a marker for lung cell senescence, in CS-exposed young p16-3MR mice **(Figure 5)**. Similarly, p53 expression was also markedly augmented in CS exposed young mice, but not in old p16-3MR mice (**Figure 6**). Though not significant, the expression of both PAI-1 and p53 were decreased in CS exposed older p16-3MR mice thus showing an age-dependent regulation of cellular senescence (**Figures 5&6**). Contrary to these, the expression of FN-1 showed slight variations in CS-exposed young and old p16-3MR mice, but none of these changes were significant **(Figure 5)**. We did not observe any change in the expression of inflammatory subunits of NF-κB (p50/p105) on sub-chronic CS exposure in both young and old p16-3MR mice (**Figure 6**).

**Figure 6:**
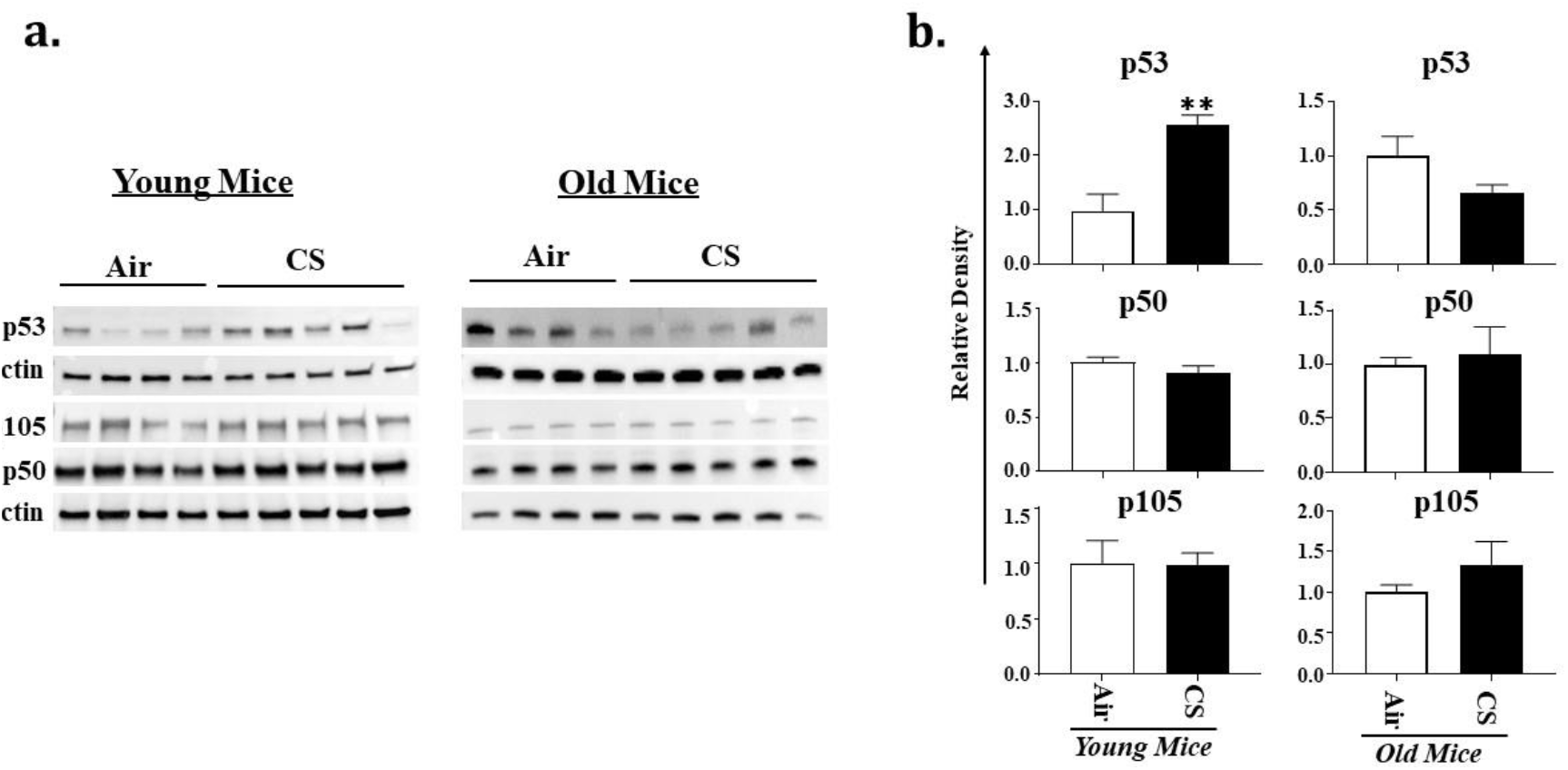
Altered expression of senescence and inflammatory marker in CS-exposed p16-3MR mice. (a) Younger and older p16-3MR mice were exposed to sub-chronic CS for 30 days and the expression of marker for cellular senescence (p53) and inflammation (p50 & p105) was determined using immunoblotting analyses. β-actin was used as loading controls. (b) The band intensity was measured by densitometry and data were shown as fold change relative to respective control group. Full Gels/blots with bands (unedited/uncropped electrophoretic gels/blots) obtained from air- and CS-exposed younger and older mouse samples from the same experiments were processed in parallel are shown (see Supplementary information as Suppl Figs. 9-10). Data are shown as mean ± SEM (n = 4–5/group). \**p* < 0.05 vs Air as per Student’s t-test for pairwise comparisons.

We further studied the mRNA expression of other SASP-related genes IL-1α, CCL2, IL-6 and CCL5 in CS-exposed young and old p16-3MR mice. Our results demonstrated age-dependent increase in the expression of IL-6 and CCL2 in CS-exposed mice as compared to the air controls. The expression of IL-1α was also significantly increased in CS-exposed young and old mice, while no change was observed in the expression of CCL5 in response to sub-chronic CS exposure (**Figure 7**).

**Figure 7:**
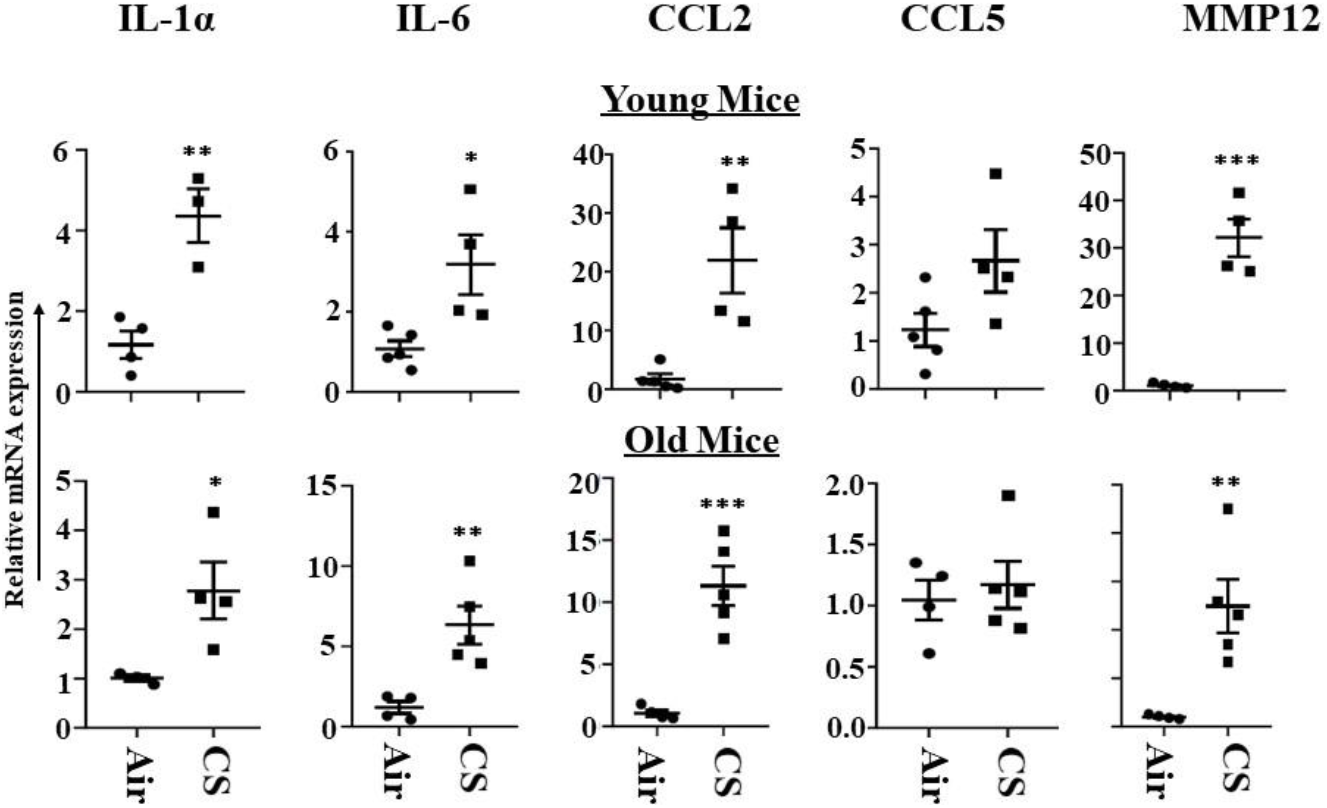
Increased mRNA expression of SASP-associated markers in CS-exposed p16-3MR mice. mRNA expression of SASP associated markers (IL-1α, IL6, CCL2, CCL5 and MMP12) were measured in the lung tissues from air- and CS-exposed young and old p16-3MR mice. Bar graphs represent the mean normalized fold change of respective air- vs. CS-exposed mouse lungs using ∆∆Ct method. Data are shown as mean ± SEM (n = 4–5/group). *p < 0.05, \*\**p* < 0.01, \*\*\**p* < 0.001 vs Air as per Student’s t-test for pairwise comparisons.

To further assess the senescence-mediated induction of pro-inflammatory markers on CS exposure (30 days), we measured the levels of pro-inflammatory cytokines/chemokines in the blood plasma of air- and CS-exposed young and old p16-3MR mice using Luminex multiplex assay. Interestingly, we observed a significant increase in the levels of eotaxin and IL-17A in CS-exposed older p16-3MR mice (**Figure 8a**).

**Figure 8:**
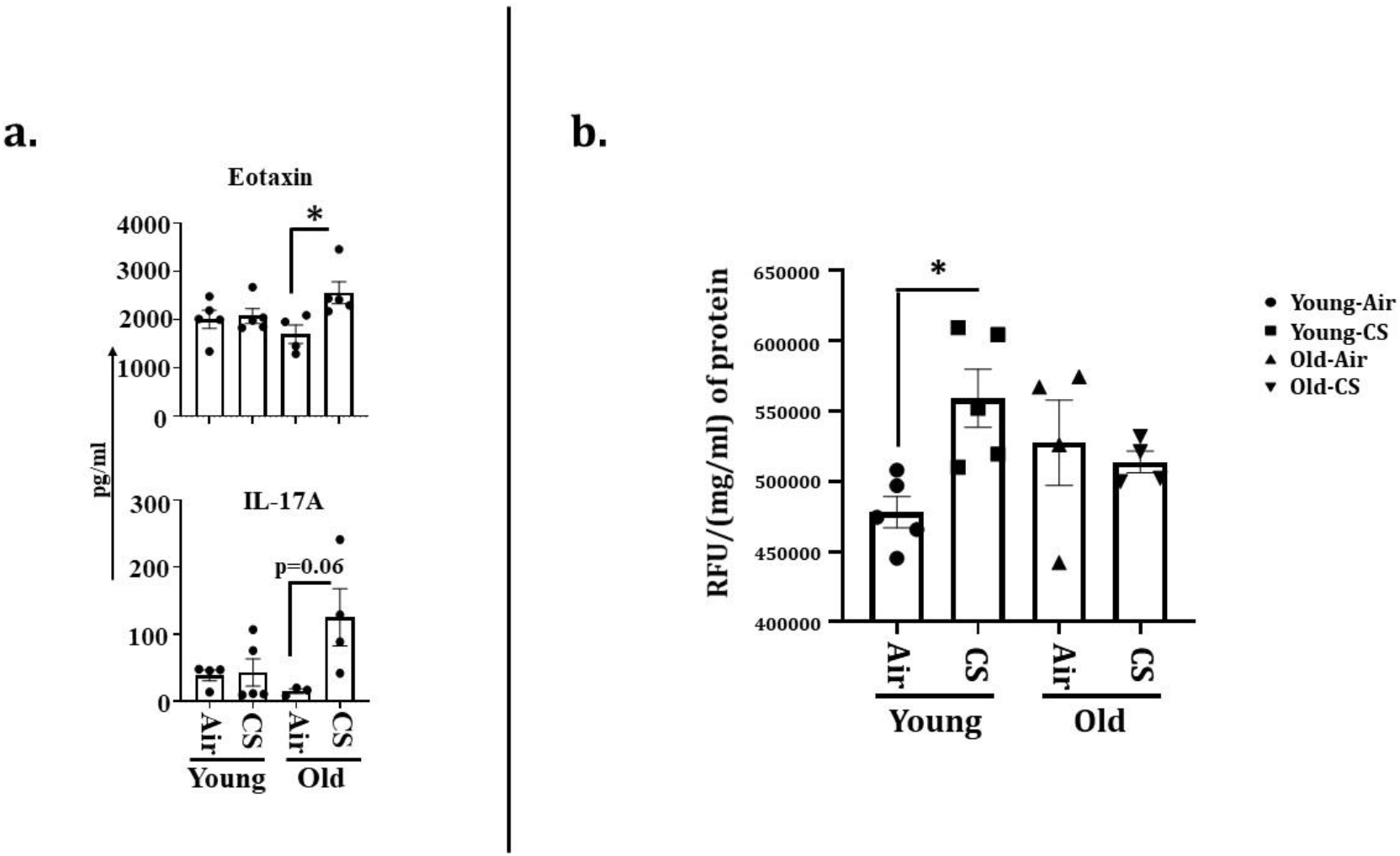
Increased level of plasma cytokines and SA-β-gal activity in CS-exposed p16-3MR mice. Younger and older p16-3MR mice were exposed to sub-chronic CS for 30 days and plasma was used to determine SASP cytokines. (**a**) The level of pro-inflammatory plasma cytokines (eotaxin and IL-17A) were measured using Luminex multiplex assay. (**b**) SA-β-gal activity was determined in lung homogenates. Data are shown as mean ± SEM (n = 4–5/group). \**p* < 0.05, \*\**p* < 0.01, \*\*\**p* < 0.001 vs Air as per Student’s t-test for pairwise comparisons.

### Sub-chronic CS exposure contributes towards the augmentation of SA-β-gal activity in the lungs of p16-3MR mice

Considering that the SA-β-gal activity (lysosomal) is a marker of cellular senescence, we measured the SA-β-gal activity in the lung homogenates of air- and CS-exposed younger and older p16-3MR mice. We observed an age-independent augmentation of SA-β-gal activity in the CS-exposed young and old mice (**Figure 8b**). Collectively, these results show that the characteristic of aging itself is involved in lung cellular senescence program/process, but does not increase the susceptibility of CS-induced cellular senescence in lung aging using a mouse model of COPD.

## DISCUSSION

Cellular senescence is a complex response towards stress where originally proliferating cells lose their power to proliferate. It is known to play an important role in two disparate processes tumorigenesis and aging. Mounting evidence suggests that the senescence-induced growth arrest acts as a barrier towards tumor growth. However, accumulation of senescent cells can also drive aging-associated pathologies. Therefore, cellular senescence is the prime example of evolutionarily antagonistic pleiotropic response, which is beneficial at young age but deleterious at older ages ^9,18,21^.

Many cigarette smoke (CS)-related pathologies show exacerbating symptoms at older age ^22–24^. Reports suggest that cellular senescence is responsible for aggravating disease symptoms in pathologies like asthma, COPD, and pulmonary fibrosis ^6,25–27^. In fact, cigarette smoke has been shown to cause DNA damage leading to premature and accelerated lung aging that ultimately leads to the development of COPD ^27,28^. However, the exact role of senescence in development of these lung pathologies is not completely understood. One of the hindrances in this regard is the lack of a suitable reporter model to study lung cellular senescence, though several models are proposed which have their own pros and cons ^19,29–32^. To overcome this hindrance, Demaria et al., (2014) generated a mouse model named p16-3MR that was shown to effectively identify, isolate and selectively eliminate senescent cells ^18^.

In this reporter mouse model, the senescence-sensitive promoter of p16^INK4a^ and its adjacent p19^Arf^ genes were inactivated and integrated in the BAC (bacteria artificial chromosome) with a 3MR transgene. This 3MR transgene encoded three fusion protein made up of luciferase (LUC), monomeric red fluorescent protein (mRFP), and Herpes Simplex Virus thymidine kinase (HSV-TK) allowing identification, sorting and selective elimination of senescent cells, respectively ^18^. In the current study, we assessed the suitability of this model to study lung cellular senescence in cigarette smoke-related pathologies. We exposed young (12-14 months) and old (17-20 months) p16-3MR mice to CS for the duration of 30 days. Thereafter, we assessed the expression of senescent markers and SASP-related genes in the lungs of control and treatment groups.

On studying the tissue luminescence in lung tissues using IVIS imaging, we found a CS-induced upregulation of p16 expression indicative of increase in tissue luminescence which was significant amongst young mice. Considering that luminescence indicates cellular senescence in this model, we were effectively able to show premature induction of senescence in CS-exposed young mice in our study.

We next studied the mRFP expression in lung tissue sections using fluorescence microscopy to track the senescent cells. Our results further substantiated our previous finding thus showing that this model could effectively be used to track the CS-induced senescent cell using fluorescence. It is pertinent to mention here that we were unable to use tissue fluorescence to identify mRFP expression in the lung tissues from air- and CS-exposed young and old p16-3MR mice. IVIS imaging results showed excessive fluorescence in the whole-lung for CS-exposed group, which could be an outcome of auto fluorescence due to tar deposition on CS inhalation in these animals. We thus demonstrated that IVIS imaging cannot be used to study senescence-induced tissue fluorescence in p16-3MR mice. However, our fluorescence microscopy and qPCR results were able to identify CS-induced senescence in our mouse model. In fact, future studies could use immunohistochemical staining to identify the specific cell types undergoing cellular senescence following chronic CS exposure.

We further confirmed CS-induced senescence by increased (a) transcript levels (mRNA) and protein expression of senescent markers p16 and p21, (b) mRNA levels of SASP factors- IL-1α, IL-6, CCL2 and MMP12, and (c) SA-β-gal activity assay in CS exposed group as compared to air controls. Our results are comparable to our previous studies ^20^, thus proving that the results obtained using this model are translatable.

Since cigarette smoke-induced emphysema results in tissue remodeling and ECM proteins we further tested the expression of matrix metalloproteinases (MMP9 and MMP12), tissue plasmin regulators (PAI-1) and ECM glycoprotein (FN-1) on 30 days CS-exposed young and old p16-3MR mice. Though not significant, our results demonstrate age-dependent dysregulation in the MMP9 protein expression in CS-exposed mice as compared to the air controls. Contrarily, we found significant increase in the levels of plasminogen activator inhibitor (PAI)-1 in CS-exposed young p16-3MR mice as compared to air-exposed control mice. Dysregulated levels of MMPs and its inhibitors has been associated with abnormal tissue repair in conditions like fibrosis and asthma ^33–35^. Thus, our results show that this model is effective in simulating conditions leading to ECM deposition and fibrinolysis on CS exposure which is similar to that found in human ^33^. Since ours was sub-chronic (30 days CS exposure) study, we could not find conditions leading to emphysematous lungs, but our current findings prove the efficacy of this model to mimic conditions of acute and chronic exposure to cigarette smoke *in vivo*.

Interestingly, we observed age-dependent changes in the expression of many proteins associated with inflammation and senescence, like, p53, IL-17A, eotaxin, on CS-exposure in our mouse model. These findings show that age plays a key role in regulating the disease phenotypes in cigarette smoke-associated disorders. In fact, the role of several proteins might vary depending on the age and duration of exposure that will be studied in the future. In the past few decades, the possibility of using senolytic drugs to combat against age-associated disorders is being considered ^15–18,36^. Given that this model can selectively target senescent cells by the use of Ganciclovir, it will be interesting how the elimination of senescence affects smoking-induced molecular changes *in vivo*. We intend to determine this possibility in our upcoming studies.

Overall, in the current study, we proved that the *in vivo* mouse model for cellular senescence using p16-3MR reporter mice could be used to study smoking-induced age-related pathologies like COPD and IPF. In general, this is the first attempt to utilize p16-3MR as a senescence model in lung pathologies (**Figure 9**). Future studies are required with a greater focus on using the p16-3MR model to discern the role of lung cellular senescence in the progression of aging and exosome-induced conditions like COPD as well as to test whether the removal of senescent cells causes any alleviation in the disease phenotype. Nevertheless, use of this model will help to deduce the cross-talk between cellular senescence and inflammation in smoking-associated pulmonary conditions to identify effective therapies in the future.

**Figure 9:**
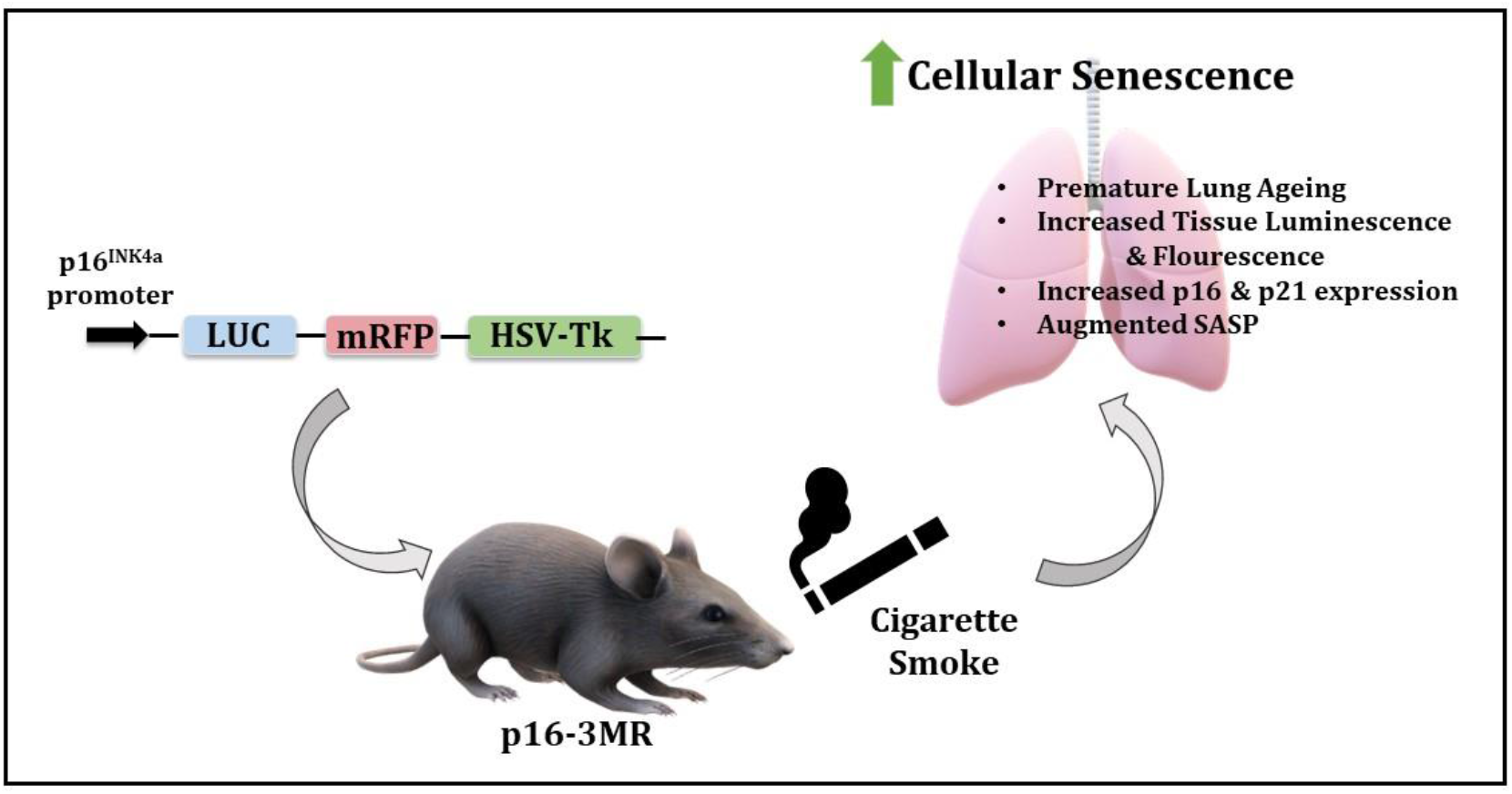
Schematics of the model and observed outcomes as shown using the p16-3MR mice exposed to cigarette smoke for cellular senescence.

## METHODS

### Ethics statement and scientific rigor/reproducibility

All animal experiments were performed according to the standards established by the U.S. Animal Welfare Act as per the NIH guidelines. The University Committee on Animal Research at the University of Rochester approved the animal protocols described below. Great precaution was taken in employing a robust and unbiased approach during the experimental and corresponding results analysis phase in order to ensure reproducibility befitting NIH standards. The key biological and chemical resources used in this study are of scientific standard and have been authenticated and revalidated. Unless stated otherwise, all biochemical reagents were purchased from Millipore Sigma (St. Louis, MO, USA). The antibodies listed in the study were commercial grade and were validated by their respective manufacturers.

### p16-3MR mouse model

We obtained the p16-3MR mice from Dr. Judith Campisi of the Buck Institute for Research on Aging via the Unity Biotechnology, Inc, San Francisco, CA for conducting our experiments. p16-3MR mice are diploid for p16^INK4a^ and p19^Arf^ with a trimodal (3MR) reporter fusion protein designed to identify, isolate, and selectively kill senescent cells ^18^. We used these mice upon genotyping to test the suitability of this model for studying cellular senescence in cigarette smoke exposure-related lung pathologies. Animals that have undergone cellular senescence show an increased expression of p16. These senesced cells could be identified with luminescence and red fluorescence protein (RFP) and selectively eliminate senescent cells by treatment with Ganciclovir in the p16-3MR mice model. All the mice were housed in the vivarium facility at the University of Rochester Medical Center with a 12-hour light/12-hour dark cycle (lights on at 6:00 am). All the animals used in the study were genotyped prior to CS exposure.

### Sub-Chronic CS exposure

Male and female mice of different age-groups (12–14 months and 17–20 months) were exposed to sub-chronic cigarette smoke generated by research grade cigarettes (3R4F) according to the Federal Trade Commission protocol (1 puff/min of 2-s duration and 35-ml volume for a total of 8 puffs at a flow rate of 1.05 L/min) with a Baumgartner-Jaeger CSM2072i automatic CS generating machine (CH Technologies, Westwood, NJ) ^37,38^. The mainstream smoke exposure was performed at a concentration of ~250–300 mg/m^3^ total particulate matter (TPM) by adjusting the flow rate of the diluted medical air, and the level of carbon monoxide in the chamber measured at ~350 ppm, as described previously ^37^. At the end of the exposure, 12–14 months old mice were referred to as “young,” whereas 17-20 months old were termed “old” mice. We will use the same terms to denote these two groups in the rest of this manuscript. Mouse that were not exposed to CS were considered the “Air” group and kept in filtered air, which served as the control group in these experiments. 24 h following the final exposure, all the mice were euthanized and their lung tissues were used for imaging, biochemical and immunohistochemical analyses.

### Tissue Luminescence and Fluorescence using IVIS Imaging

To identify senescent cells using luminescence, lung tissues harvested from euthanized mice were soaked for 10 min in pre-warmed (37°C) PBS with 2% FBS and a 1:10 dilution of Xenolight RediJect Coelenterazine h (Cat# 706506, Perkin Elmer, Waltham, MA). Following a 12-15 min incubation, tissues were transferred to a fresh 35-mm dish and luminescence was measured using the IVIS® Spectrum multispectral imaging instrument (Caliper Life Sciences, Inc. - Hopkinton, MA). IVIS® Spectrum multispectral imaging instrument was also used to measure the lung tissue fluorescence (RFP) in our control and experimental groups at an excitation maximum of 535 nm and emission maximum of 580 nm.

### Fluorescence microscopy in Lung Tissue section

Fluorescence imaging was employed to determine mRFP expression in the lung tissue sections from CS- and air-exposed mice. More specifically, non-lavaged mouse lungs were inflated with 50% solution of Optimal Cutting Temperature (OCT) compound and 10 μm frozen sections were cut using a rotary microtome-cryostat. Immediately after sectioning, samples were fixed and mounted using Prolong (Cat# P36962, Life technologies Corporation, Eugene, OR) with DAPI. Images were acquired using Nikon Eclipse Ni-U fluorescence microscope at 200× magnification using Advance SPOT software.

### mRNA expression analyses using qPCR

Reverse-Transcriptase Polymerase Chain Reaction (RT-PCR) was performed to determine the differential expression of cellular senescence genes in our experimental groups. Briefly, RNA was isolated from the lung tissue using RNeasy miniprep kit (Qiagen, Valencia, CA, USA). RNA quantity and quality were assessed using Nanodrop 1000 spectrophotometer (Thermo Fisher Scientific) and 1 μg of total RNA was used for cDNA conversion using the RT2 first strand kit (Cat# 330404, Qiagen, Valencia, CA, USA). The prepared cDNA was diluted and used to determine the expression of genes of interest using specific primers. All of the real-time PCR reactions were performed with RT2 SYBR Green/ROX PCR Master Mix (Cat# 330503, Qiagen, Valencia, CA, USA) and relative mRNA expression of each gene was determined using CFX96 real-time system (Bio-Rad, Hercules, CA). Differential expression of target genes in total RNA isolated from air- and CS-exposed lung tissues from young and old p16-3MR mice were expressed as relative fold change. 18S rRNA was used as housekeeping control. Fold-Change (2^(- Delta Delta Ct)) is the normalized gene expression (2^(- Delta Ct)) in the CS exposed group (Treated) divided by the normalized gene expression (2^(- Delta Ct)) in the Air group (Control) ^39,40^. The sequence of the primers used for amplification is provided in Supplementary file Table 1.

### Immunoblot Analysis

To determine the protein expression of various senescence-associated proteins, we employed immunoblotting. Briefly, one lobe of the lung tissue (~40 mg) was homogenized (Pro 200 homogenizer, at maximum speed, 5th gear for 40 seconds) in 0.3 mL of ice-cold RIPA buffer containing complete protease inhibitor cocktail (Cat# 78444, Thermo Scientific, Waltham, MA). The tissue homogenate was incubated on ice for one hour to allow complete cell lysis. Following this incubation, the homogenate was centrifuged at 13,000*g* for 30 minutes at 4°C. The supernatant was aliquoted and stored at −80°C until further analyses. A small fraction of the tissue lysate was taken and diluted for protein analysis with the help of bicinchoninic acid (BCA) colorimetric assay (Thermo Scientific, Waltham, MA) where BSA was used as a standard.

Equal amounts of protein from each sample (20 μg) was resolved on a sodium dodecyl sulfate (SDS)-polyacrylamide gel and subsequently electroblotted onto nitrocellulose membranes. The nitrocellulose membranes were then blocked using 5% milk solution for one hour at room temperature. Thereafter, the membranes were probed with antibodies targeted for the protein of interest. Antibodies towards anti-p16 (Cat# sc-377412, Santa Cruz Biotechnology, Dallas, TX), anti-p21 (Cat# 556430, BD Biosciences, San Jose, CA), anti-Fn1(Cat# ab2413), anti-PAI-1 (Cat# ab66705), anti-NF-κB(Cat# ab32360), anti-MMP9 (Cat# ab38898) (Abcam, Cambridge, UK), anti-MMP12 (Cat# NBP2-67344, Novus Biologicals, Littleton, CO), anti-p53 (Cat# 2524, Cell Signaling, Danvers, MA) and anti-β-actin (Cat# 12620, Cell Signaling, Danvers, MA) were added using a 1:1000 dilution (in 5% BSA in tris-buffered saline [TBS] containing 0.1% Tween 20) and incubated overnight at 4°C. The following day, blots were washed three times with 1X TBST and probed with the appropriate anti-rabbit (Cat# 170-6515, Bio-Rad, Hercules, CA) or anti-mouse (Cat# 7076, Cell Signaling, Danvers, MA) secondary antibody (1:3000 dilution in 5% milk) for one hour. Chemiluminescence was detected through the Bio-Rad ChemiDoc MP Imaging system using the SuperSignal West Femto Maximum Sensitivity Substrate (Cat# 34096, Thermo Scientific, Waltham, MA). Bio-Rad Image Lab software was used for densitometric analyses. Corresponding band intensities for the proteins of interest were plotted as fold change relative to their respective β-actin loading control bands.

### Measurement of SA-β-gal activity

SA-β-gal activity was measured using cellular senescence activity assay kit (Cat# ENZ-KIT129-0120, Enzo Life sciences, Farmingdale, NY) as per the manufacturer’s protocol. Briefly, one lobe of the lung tissue (~40 mg) was homogenized (Pro 200 homogenizer) in 0.3 mL of ice-cold 1X Cell lysis buffer containing complete protease inhibitor cocktail (Cat# 78444, Thermo Scientific, Waltham, MA). Tissue homogenate was incubated on ice for 30 min and then centrifuged at 13,000 rpm for 15 min at 4°C. The supernatant was collected and stored until further analyses. 50 ml of cell lysate was mixed with 50 ul of assay buffer and incubated for 3 hours at room temperature. Following incubation, 50 ul of the reaction mixture was added to 200 ul of Stop solution and fluorescence was read using Cytation 5 (Biotek, Winooski, VA) at 360 nm (Excitation) / 465 nm (Emission).

### Assessment of pro-inflammatory mediators using Luminex

The level of proinflammatory mediators in plasma (50 μL) was measured with the help of Bio-Plex Pro Mouse Cytokine Standard 23-Plex (Cat# 64209360, Bio-Rad, Hercules, CA) as per manufacturer’s protocol. Blood plasma was diluted two-folds and the levels of each cytokine/chemokine were expressed as pg/ml.

### Statistical analysis

All statistical calculations were performed using GraphPad Prism 8.0. Data are expressed as mean ± SEM. Pairwise comparisons were done using unpaired Student’s *t*-test. For multi-group comparisons, one-way Analysis of Variance (ANOVA) with ad-hoc Tukey’s test was employed. All animal experiments (n=4–5 mice/group) were performed twice. Differences were considered statistically significant at \**p* < 0.05, \*\**p <* 0.01, and \*\*\**p <* 0.001 when compared with respective air controls.

## Acknowledgments

The National Institutes of Health (NIH) 1R01HL135613 R01 ES029177, HL137738 and R01 HL133404 supported this study. The funding body has no role in design of the study, data collection, analysis, and interpretation of data and in writing the manuscript. We thank Dr. Judith Campisi of the Buck Institute for Research on Aging via the Unity Biotechnology, Inc, San Francisco, CA for providing the p16-3MR strain for our research purposes. We thank Mr Shaiesh Yogeshwaran for his technical and editing help.

## Authors Contributions

GK and ISK designed and conducted the experiments; IR conceived the concept and ideas; GK, ISK, and IR wrote and/or edited/revised the manuscript. IR obtained research funding.

## Declarations

**Disclosures: Conflict of or competing interest statement**

The authors have declared that no competing interests exist.

## References

1 Tuder, R. M., Kern, J. A. & Miller, Y. E. Senescence in chronic obstructive pulmonary disease. Proceedings of the American Thoracic Society 9, 62–63, doi:10.1513/pats.201201-012MS (2012).

2 Karrasch, S., Holz, O. & Jörres, R. A. Aging and induced senescence as factors in the pathogenesis of lung emphysema. Respiratory medicine 102, 1215–1230, doi:10.1016/j.rmed.2008.04.013 (2008).

3 Sundar, I. K., Rashid, K., Gerloff, J., Li, D. & Rahman, I. Genetic Ablation of p16(INK4a) Does Not Protect against Cellular Senescence in Mouse Models of Chronic Obstructive Pulmonary Disease/Emphysema. American journal of respiratory cell and molecular biology 59, 189–199, doi:10.1165/rcmb.2017-0390OC (2018).

4 Lee, J., Sandford, A., Man, P. & Sin, D. D. Is the aging process accelerated in chronic obstructive pulmonary disease? Curr Opin Pulm Med 17, 90–97, doi:10.1097/mcp.0b013e328341cead (2011).

5 Zhou, F., Onizawa, S., Nagai, A. & Aoshiba, K. Epithelial cell senescence impairs repair process and exacerbates inflammation after airway injury. Respiratory research 12, 78, doi:10.1186/1465-9921-12-78 (2011).

6 Tsuji, T., Aoshiba, K. & Nagai, A. Alveolar cell senescence in patients with pulmonary emphysema. American journal of respiratory and critical care medicine 174, 886–893, doi:10.1164/rccm.200509-1374OC (2006).

7 Nyunoya, T. et al. Cigarette smoke induces cellular senescence. American journal of respiratory cell and molecular biology 35, 681–688, doi:10.1165/rcmb.2006-0169OC (2006).

8 Amsellem, V. et al. Telomere dysfunction causes sustained inflammation in chronic obstructive pulmonary disease. American journal of respiratory and critical care medicine 184, 1358–1366, doi:10.1164/rccm.201105-0802OC (2011).

9 Campisi, J. & d’Adda di Fagagna, F. Cellular senescence: when bad things happen to good cells. Nature Reviews Molecular Cell Biology 8, 729–740, doi:10.1038/nrm2233 (2007).

10 Wallis, R., Mizen, H. & Bishop, C. L. The bright and dark side of extracellular vesicles in the senescence-associated secretory phenotype. Mechanisms of ageing and development 189, 111263, doi:10.1016/j.mad.2020.111263 (2020).

11 Franceschi, C. & Campisi, J. Chronic inflammation (inflammaging) and its potential contribution to age-associated diseases. The journals of gerontology. Series A, Biological sciences and medical sciences 69 Suppl 1, S4–9, doi:10.1093/gerona/glu057 (2014).

12 Liu, R. M. & Liu, G. Cell senescence and fibrotic lung diseases. Experimental gerontology 132, 110836, doi:10.1016/j.exger.2020.110836 (2020).

13 van Deursen, J. M. The role of senescent cells in ageing. Nature 509, 439–446, doi:10.1038/nature13193 (2014).

14 Childs, B. G. et al. Senescent cells: an emerging target for diseases of ageing. Nat Rev Drug Discov 16, 718–735, doi:10.1038/nrd.2017.116 (2017).

15 Patil, P. et al. Systemic clearance of p16(INK4a) -positive senescent cells mitigates age-associated intervertebral disc degeneration. Aging cell 18, e12927, doi:10.1111/acel.12927 (2019).

16 Kim, H.-N. et al. Elimination of senescent osteoclast progenitors has no effect on the age-associated loss of bone mass in mice. Aging cell 18, e12923–e12923, doi:10.1111/acel.12923 (2019).

17 Rocha, L. R. et al. Early removal of senescent cells protects retinal ganglion cells loss in experimental ocular hypertension. Aging cell 19, e13089, doi:10.1111/acel.13089 (2020).

18 Demaria, M. et al. An essential role for senescent cells in optimal wound healing through secretion of PDGF-AA. Dev Cell 31, 722–733, doi:10.1016/j.devcel.2014.11.012 (2014).

19 Liu, J.-Y. et al. Cells exhibiting strong p16(INK4a) promoter activation in vivo display features of senescence. PNAS 116, 2603–2611, doi:10.1073/pnas.1818313116 %J Proceedings of the National Academy of Sciences (2019).

20 Rashid, K., Sundar, I. K., Gerloff, J., Li, D. & Rahman, I. Lung cellular senescence is independent of aging in a mouse model of COPD/emphysema. Scientific reports 8, 9023, doi:10.1038/s41598-018-27209-3 (2018).

21 Paez-Ribes, M., González-Gualda, E., Doherty, G. J. & Muñoz-Espín, D. Targeting senescent cells in translational medicine. EMBO molecular medicine 11, e10234, doi:10.15252/emmm.201810234 (2019).

22 Ito, K. & Barnes, P. J. COPD as a disease of accelerated lung aging. Chest 135, 173–180, doi:10.1378/chest.08-1419 (2009).

23 Selman, M., López-Otín, C. & Pardo, A. Age-driven developmental drift in the pathogenesis of idiopathic pulmonary fibrosis. The European respiratory journal 48, 538–552, doi:10.1183/13993003.00398-2016 (2016).

24 Nash, S. H., Liao, L. M., Harris, T. B. & Freedman, N. D. Cigarette Smoking and Mortality in Adults Aged 70 Years and Older: Results From the NIH-AARP Cohort. Am J Prev Med 52, 276–283, doi:10.1016/j.amepre.2016.09.036 (2017).

25 Schafer, M. J. et al. Cellular senescence mediates fibrotic pulmonary disease. Nature communications 8, 14532, doi:10.1038/ncomms14532 (2017).

26 Wu, J. et al. Central role of cellular senescence in TSLP-induced airway remodeling in asthma. PLoS One 8, e77795, doi:10.1371/journal.pone.0077795 (2013).

27 Brandsma, C. A. et al. Lung ageing and COPD: is there a role for ageing in abnormal tissue repair? European respiratory review : an official journal of the European Respiratory Society 26, doi:10.1183/16000617.0073-2017 (2017).

28 Durham, A. L. & Adcock, I. M. The relationship between COPD and lung cancer. Lung Cancer 90, 121–127, doi:10.1016/j.lungcan.2015.08.017 (2015).

29 Sorrentino, J. A. et al. p16INK4a reporter mice reveal age-promoting effects of environmental toxicants. The Journal of clinical investigation 124, 169–173, doi:10.1172/jci70960 (2014).

30 Burd, Christin E. et al. Monitoring Tumorigenesis and Senescence In Vivo with a p16INK4a-Luciferase Model. Cell 152, 340–351, doi:https://doi.org/10.1016/j.cell.2012.12.010 (2013).

31 Omori, S. et al. Generation of a p16 Reporter Mouse and Its Use to Characterize and Target p16high Cells In Vivo. Cell Metabolism 32, 814–828.e816, doi:https://doi.org/10.1016/j.cmet.2020.09.006 (2020).

32 Folgueras, A. R., Freitas-Rodríguez, S., Velasco, G. & López-Otín, C. Mouse Models to Disentangle the Hallmarks of Human Aging. Circulation research 123, 905–924, doi:10.1161/circresaha.118.312204 (2018).

33 Oh, C. K., Ariue, B., Alban, R. F., Shaw, B. & Cho, S. H. PAI-1 promotes extracellular matrix deposition in the airways of a murine asthma model. Biochemical and biophysical research communications 294, 1155–1160, doi:https://doi.org/10.1016/S0006-291X(02)00577-6 (2002).

34 Freitas-Rodríguez, S., Folgueras, A. R. & López-Otín, C. The role of matrix metalloproteinases in aging: Tissue remodeling and beyond. Biochimica et Biophysica Acta (BBA) - Molecular Cell Research 1864, 2015–2025, doi:https://doi.org/10.1016/j.bbamcr.2017.05.007 (2017).

35 Atkinson, J. J. et al. The role of matrix metalloproteinase-9 in cigarette smoke-induced emphysema. American journal of respiratory and critical care medicine 183, 876–884, doi:10.1164/rccm.201005-0718OC (2011).

36 Kirkland, J. L. & Tchkonia, T. Senolytic drugs: from discovery to translation. J Intern Med 288, 518–536, doi:10.1111/joim.13141 (2020).

37 Yao, H. et al. Cigarette smoke-mediated inflammatory and oxidative responses are strain-dependent in mice. Am J Physiol Lung Cell Mol Physiol 294, L1174–1186, doi:10.1152/ajplung.00439.2007 (2008).

38 Rajendrasozhan, S., Chung, S., Sundar, I. K., Yao, H. & Rahman, I. Targeted disruption of NF-{kappa}B1 (p50) augments cigarette smoke-induced lung inflammation and emphysema in mice: a critical role of p50 in chromatin remodeling. Am J Physiol Lung Cell Mol Physiol 298, L197–209, doi:10.1152/ajplung.00265.2009 (2010).

39 Sundar, I. K. & Rahman, I. Gene expression profiling of epigenetic chromatin modification enzymes and histone marks by cigarette smoke: implications for COPD and lung cancer. Am J Physiol Lung Cell Mol Physiol 311, L1245–L1258, doi:10.1152/ajplung.00253.2016 (2016).

40 Sundar, I. K. et al. Genetic ablation of histone deacetylase 2 leads to lung cellular senescence and lymphoid follicle formation in COPD/emphysema. FASEB J 32, 4955–4971, doi:10.1096/fj.201701518R (2018).

